# Colonization of *Anopheles coustani*, a neglected malaria vector in Madagascar

**DOI:** 10.1101/2024.04.16.589755

**Authors:** Tsarasoa M. Andrianinarivomanana, Fenomiaranjara T. Randrianaivo, Mandaniaina R. Andriamiarimanana, Mihary R. Razafimamonjy, Haja J.S Velonirina, Nicolas Puchot, Romain Girod, Catherine Bourgouin

**Affiliations:** Institut Pasteur de Madagascar, Medical Entomology Unit, Antananarivo, Madagascar; Institut Pasteur, Université de Paris Cité, Biology of Host-Parasite Interaction des Interactions Hôte-parasite, Paris, France

**Keywords:** *Anopheles coustani*, colony, forced mating, rearing, Madagascar

## Abstract

*Anopheles coustani* has long be recognized as a secondary malaria vector in Africa. It has recently been involved in the transmission of both *Plasmodium falciparum* and *Plasmodium vivax* in Madagascar. As most secondary malaria vector, *An. coustani* is mainly biting outdoor, which renders the control of this mosquito species difficult by the classical malaria control measures as the use of bed nets or indoor residual spraying of insecticides. The absence of a colony hinders a better understanding of its biology and vector competence towards the development of adapted mosquito control strategies. Here, we report the first successful establishment of an *An. coustani* colony from mosquito collected in Madagascar. We used a forced copulation procedure as this mosquito species will not mate in cages. We describe our mosquito colonization procedure with detailed biological features as larval to adult development and survival, recorded over the first six critical generations. The procedure should be easily applicable to *An. coustani* from different African countries, facilitating local investigation on *An. coustani* vector competence and insecticide resistance using the colony as a reference.

## Introduction

Malaria is still a public health concern in many countries, mainly in Africa [38]. For countries close to malaria elimination residual malaria is of high concern. In this regard, the role of the so-called secondary malaria vectors, often outdoor biters, in maintaining malaria transmission has been considered [39]. However, for decades most laboratory and field studies focused on the major or dominant malaria vectors [34].

With the stagnation in worldwide malaria incidence over the last decade, it is more and more important to better understand the bionomics of secondary malaria vectors for which limited data are available. This is especially crucial for secondary malaria vectors in Madagascar, a country that records a worrying increase in malaria cases despite indoor residual spraying and insecticidal treated net distribution campaigns (WHO 2022).

*Anopheles coustani* is an Afrotropical anopheline species belonging to the *Anopheles* subgenus. In several African countries including Madagascar *An. coustani* is now widely recognized as a secondary vector of human malaria parasites [15, 17, 20, 24, 26]. Although it was reported both anthropophilic and exophagic (feeding outdoors on humans) in South Africa [10], *An. coustani* is often referred as a zoophilic mosquito [35]. Nevertheless, because of its abundance it can be a locally major malaria vector [15, 28]. Indeed, in Madagascar, *An. coustani* was shown to be the major vector of both *Plasmodium vivax* and *Plasmodium falciparum* in a village where it showed a high anthropophilic behavior, associated to both endophilic and exophilic behavior [18]. Furthermore, *An. coustani* has been implicated as a potential vector of Rift Valley Fever virus in Madagascar [31] and of Chikungunya virus in Senegal [13].

*An. coustani* Laveran was first described in 1900 by Alphonse Laveran from samples received from Madagascar [23]. *Anopheles mauritianus* described at about the same time from mosquitoes collected in the Mascarene Islands was later considered as identical to *An. coustani* Laveran [[14, 32], details in [11]]. Using cytogenetics on larval polytene chromosomes, M. Coetzee reported the existence of two cryptic species within *An. coustani* collected in South Africa and Eswatini (former Swaziland) [10]. When crossed, these species named species A and species B produced infertile male F1. More recently, using *An. coustani* samples from Madagascar and morphology as well as cytogenetics, the same author confirmed that the Malagasy mosquitoes belong to the former species A, and conclude that species B was a different species named *An. crypticus* [11]. Of note, Malagasy specimens were then referred as neotypes as the original type specimens were lost. In 2020, sampling the genetic diversity of *An. coustani* population in Zambia and the Democratic Republic of Congo revealed two distinct phylogenetic groups [8].

To our knowledge no colony of *An. coustani* has ever been established. As for other species, having at hand a colony will be instrumental for various types of studies: insecticide assays with the colony as a reference, behavioral study for refining vector control strategies, assessing whether the bacteria Asaia or Wolbachia could be considered as additional tools for controlling mosquito population or their vectorial competence toward *Plasmodium* and viruses [6, 7, 33].

Establishing a novel mosquito colony from field collected samples can be very tricky. The main challenge is mating behavior. Indeed, many anopheline mosquitoes are eurygamous, mating in open space often in swarm, and anopheline females are also known for their refractoriness to multiple mating [9, 36]. *An. coustani* is no exception, as evidenced in preliminary assays while working with large number of F1 progeny obtained by in-tube forced oviposition of gravid wild females [27]. Even if F1 females would readily feed on blood, none would lay eggs over more than a week post blood feeding. Observation of a subset of the spermatheca of F1 females revealed that none had been fertilized (Nepomichene and Bourgouin, unpublished).

To overcome the absence of mating in a constrained environment (rearing cages), a «forced mating” technology was developed [4], and successfully used for establishing anopheline colonies [1, 29] even if the technique is highly demanding in human resources and skill. However, forced mating does not equal female fertilization. Indeed, it was reported that males from major anopheline vectors from the *Nyssorhynchus* subgenus including *Anopheles darlingi* would readily copulate by the forced mating technique but would fail to inseminate the females ([22]; Puchot, personal observation). On the contrary, several colonies of anophelines from both the *Anopheles* and *Cellia* subgenus were created and maintained using the forced mating technique [1, 3, 29].

Based on successful published records on anopheline from the *Anopheles* subgenus and our preliminary observations, we initiated the establishment of the first *An. coustani* colony using the forced mating technique. Here, we provide details on the major initial steps till the obtention of the eighth generation and discuss ways of improvement toward the establishment of a free-mating colony of *An. coustani*.

## Material and Methods

### Producing the F1 generation from wild *An. coustani*

*An. coustani* females were captured in Maroharona (17°36’44.33”S, 46°56’2.11”E), municipality of Andriba (Maevatanana district) in the Northwest fringe of the Central Highlands of Madagascar, in September 2019. The mosquitoes were collected in zebu parks using mouth aspirators as previously described [27]. Engorged females were transferred into 30 cm^3^ cages, provided with a 10% sucrose solution on cotton bowls. The cages were covered with a wet black fabric and transported over the next day to the Medical Entomology Unit at the Institut Pasteur de Madagascar in Antananarivo, located 6h drive away from Andriba. Morphological identification on lived mosquitoes was confirmed one by one using a dichotomous identification key adapted from [12, 16, 19]. Females were transferred to a rearing cage (32.5 x 32.5 x 32.5cm BugDorm-4® -Bioquip, Rancho Dominguez, CA, USA). A beaker half-filled with dechlorinated water was placed inside the cage for egg collection and surveyed every day for 5 days. Each batch of eggs was treated 10 min with a 0.01% formaldehyde solution and extensively rinsed with dechlorinated water for limiting potential contamination with micro-organisms as microsporidia. Eggs were then transferred to larval rearing pans (26 x 26 x 11 cm (Gilac®), filled with 1L of dechlorinated tap water (water height ∼1.5 cm). L1 and L2 larvae were fed on Tetramin Baby fish food (Tetra®) whereas L3 and L4 were fed on ground cat food. When nymphs started appearing, they were counted every day and transferred to a small beaker containing dechlorinated water and placed in a novel rearing cage. A 10% sucrose solution containing cotton bowl was placed on top of the rearing cage and changed every other day.

### Insectary conditions

Larvae and adults were reared at 26.9±0.7°C, 77±7.7% RH, with a 12:12 night and day light regimen.

### Establishing the *An. coustani* colony

From the F1 adults obtained from wild F0 mosquitoes, the artificial mating (forced mating also called forced copulation) strategy of males and females was used. This followed the procedure initially described by Baker and coll. [4], refined by Ow Yang and coll. [29] and nicely documented in [25]. Females were given a blood meal 24h before forced copulation. Males and females were cold anesthetized just before the process as illustrated in Figure 1 panels B&C. Each male (3-5 day-old) was used for fertilizing 2 females (5-10 day-old). We found unnecessary to decapitate males for copulation stimulation. After forced copulation, females were transferred to a novel rearing cage and provided a blood meal every other day to stimulate egg production. From egg production to adult emergence, the procedure described above was used and led to the successfully production of eight generations between September 2019 and May 2020. Because of the general movement restriction imposed in Madagascar during the Covid-19 outbreak, the colony could not be maintained further than the F8.

**Figure 1:**
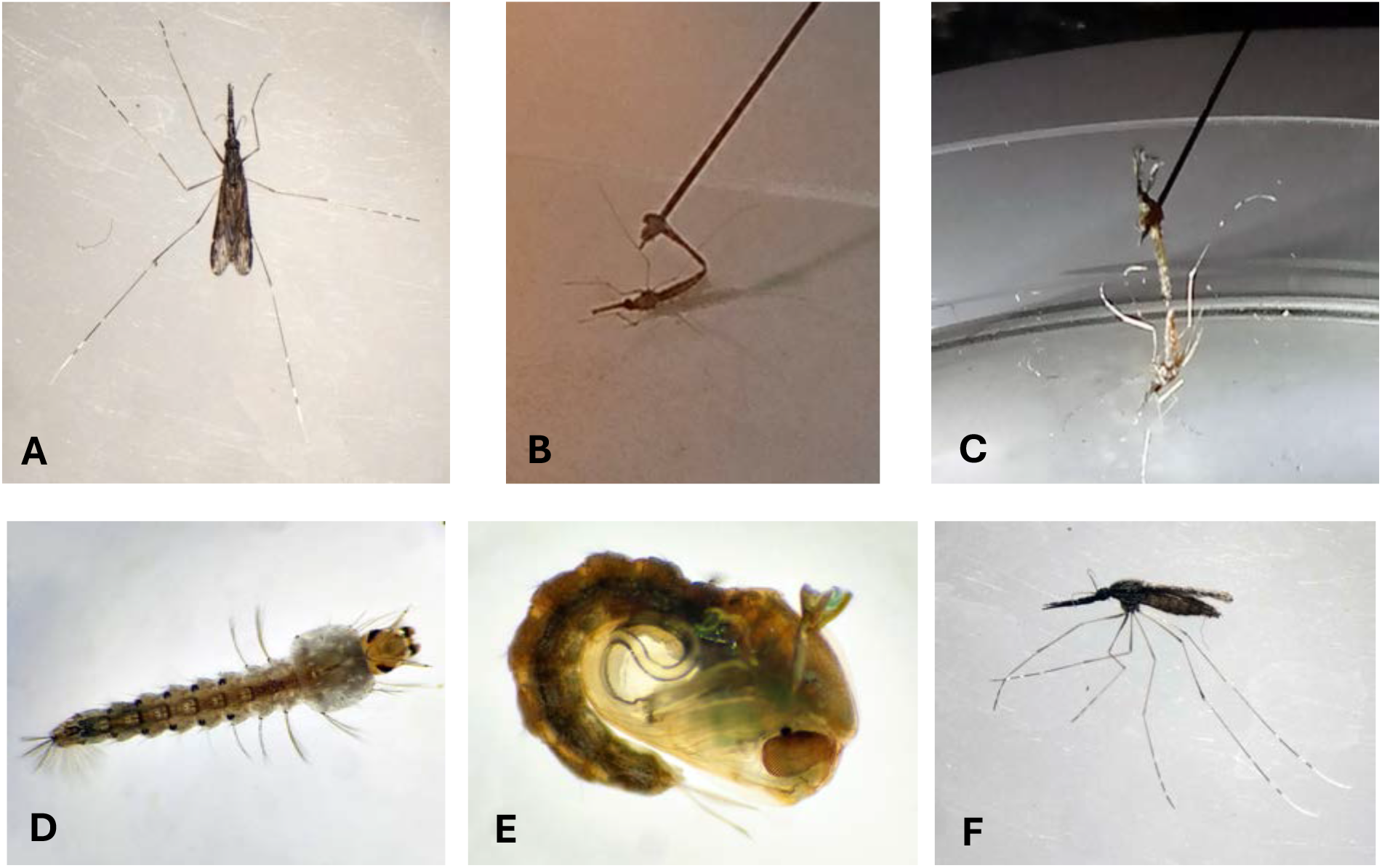
Illustrations of *An. coustani* forced copulation procedure and developmental stages. Female *An. coustani* (A and F). Forced copulation: the male is hooked on a dissecting needle and presented to a cold anesthetized female; decapitated male (B) ; non-decapitated male (C). Fourth instar larvae (D). Nymph (E).

### Recorded rearing parameters

At each generation, the following parameters were recorded: number of females successfully fertilized and their egg production, L1 to pupae growing success, as well as larvae to adult developmental time.

## Results and Discussion

### Establishing the first six generations of *An. coustani*

Starting with 31 gravid F0 wild females free to lay eggs in cage, we obtained 260 eggs, which is quite a low and challenging number (8.4 eggs/ female). This number is actually very similar to previous results [27]. The 260 eggs were obtained from a unique gonotrophic cycle of the females as they refused to feed on blood provided through the Hemotek membrane feeding device. For the sake of the project, the females from each subsequent generation were fed on rabbit a multiple times.

As previous attempts revealed that *An. coustani* would not mate in cages, the production of each generation was performed by artificial copulation at the ratio of 1 male for 2 females. It was not necessary to decapitate the male to obtain an efficient copulation and insemination of the females. Table 1 summarizes the number of males and females used for forced mating at each generation as well as the number of blood meals that the mated females received, and the mean number of eggs laid per female at each generation. The mean number of eggs produced per female increased slightly between F1 and F5, despite a reduced number of blood meals. This is possibly due to a better mastering of the forced copulation technique. Accordingly, this led to an increase of total adults from F1 to F6 (Figure 2). The production of F7 and F8 was achieved, however no data on egg and adult numbers was recorded due to the Covid-19 lockdown.

**Table 1:**
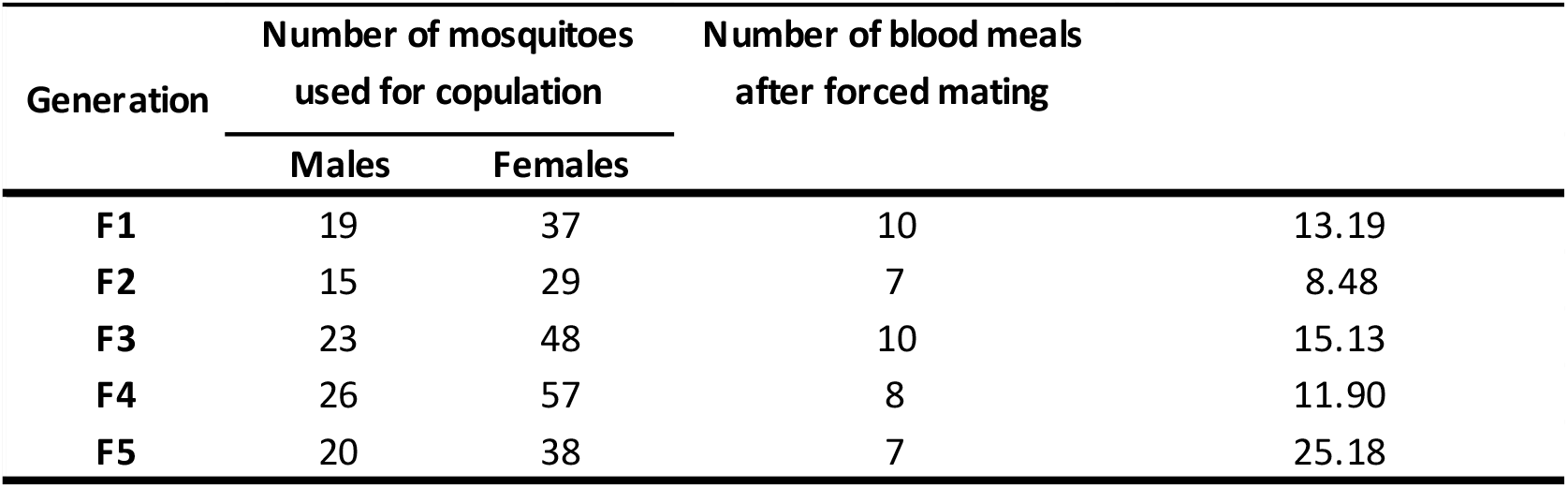
Number of males and females used at each generation in the forced mating procedure and mean egg production by each mated female.

**Figure 2:**
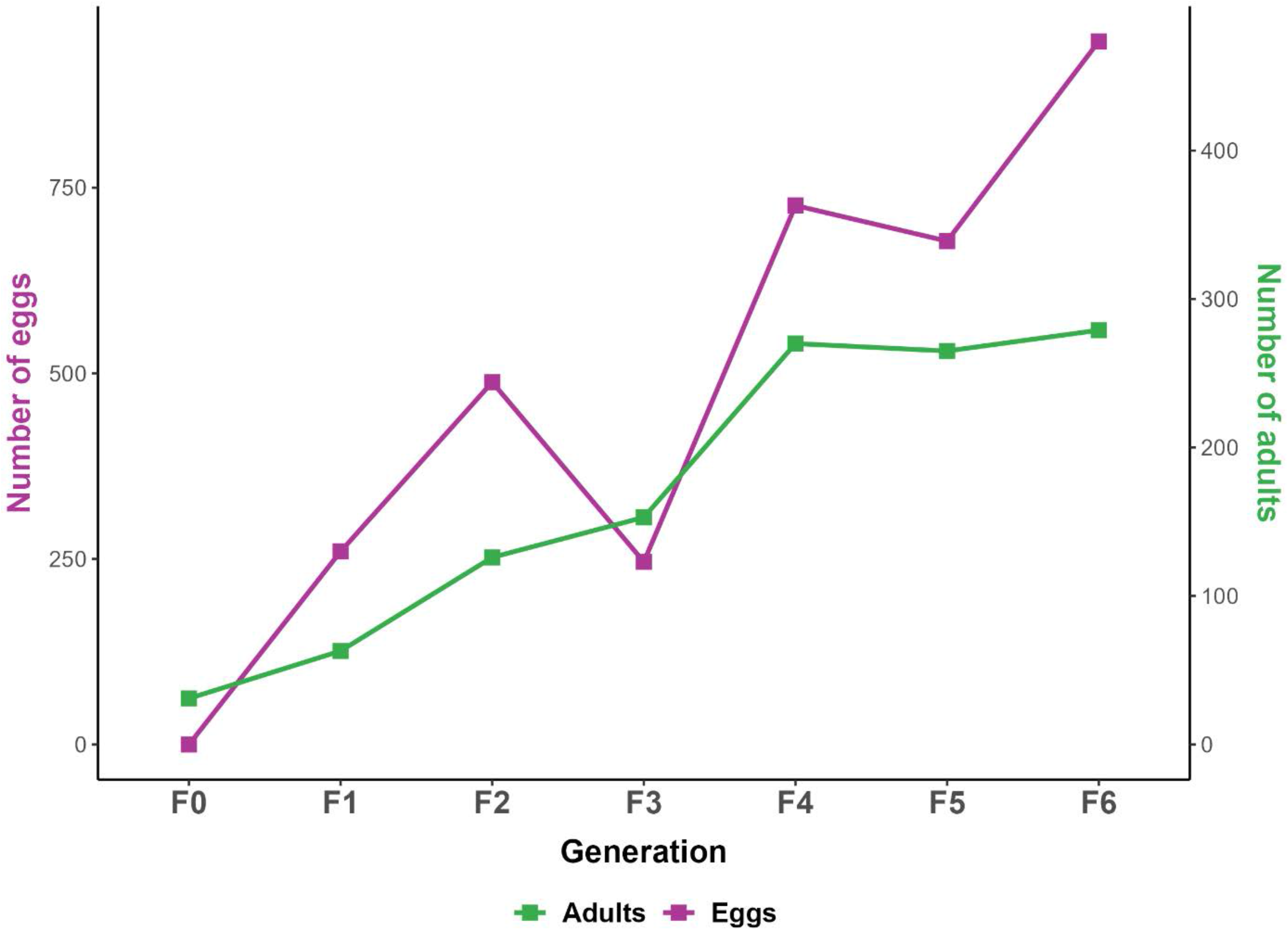
Yield in eggs and adults from F1 to F6.The graph represent the number of eggs obtained at each generation (Purple line and left scale) and the subsequent number of adults (green line and right scale).

### Bionomic features

#### Egg to adult production

As depicted in Table 2, each generation suffered a drastic reduction, with a mean survival rate from egg to adult of 36.3 % with a broad range from 24.2 to 62.2%. This latter value happened at generation F3, but no clue of this result could be identified. Nevertheless, egg to adult survival rate increased slightly across the generation and was higher than previous results on F1 only [21]. Except for the F3 generation a high mortality rate occurred during the larval stages. Normalisation of larval density and food amount might contribute to limit this rather high mortality rate. It is also striking that the mortality rate of nymphs was quite high (18.7 to 44.7 %). Such mortality might be a consequence of poor larval development linked to restricted food availability and larval competition.

**Table 2:**
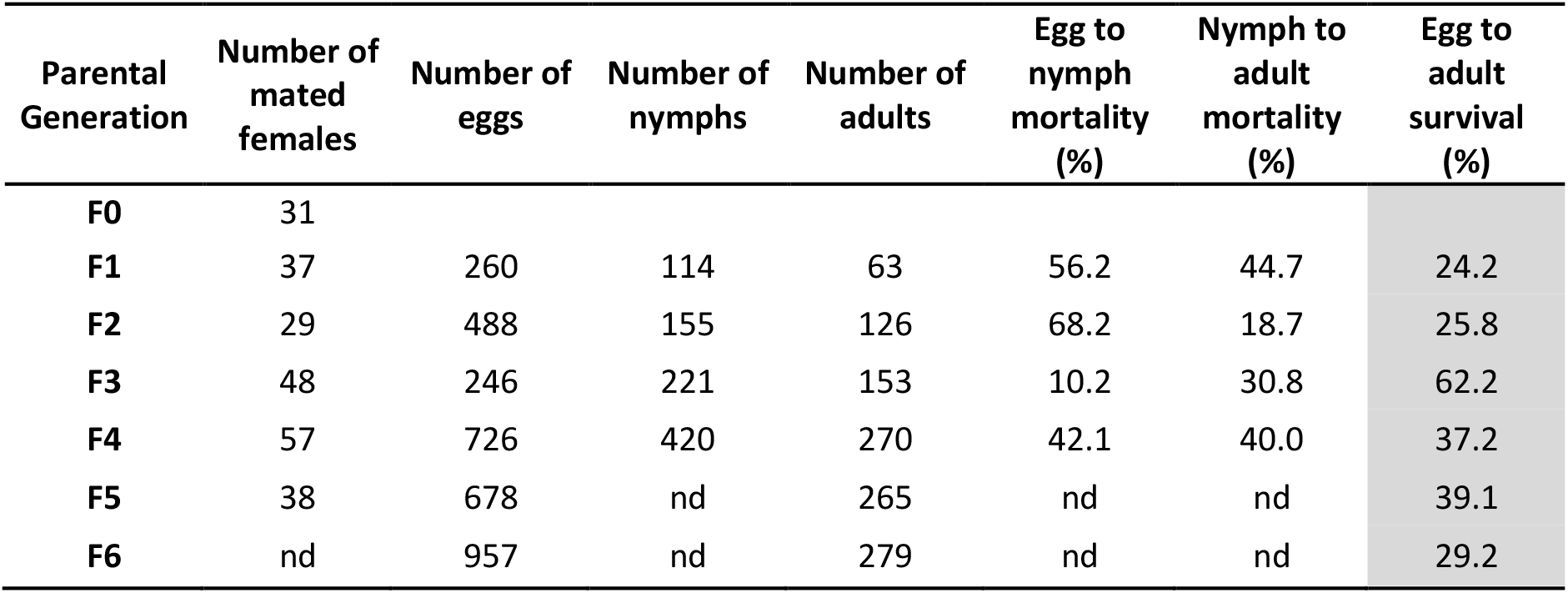
Summary of the raw data obtained across the production of the first six generation of *An. Coustani* starting with 31 wild gravid females producing 260 eggs.

#### Larval to adult development

Previous work in a field station (Andriba, Madagascar) revealed that the larval to adult development of *An. coustani* F1 was quite asynchronous and long. Indeed, the larval development lengthened more than 25 days, while *An. arabiensis* F1 population produced side by side would reached adult stage in 10 to 15 days as usually observed in insectary with tightly controlled parameters (Bourgouin et al., unpublished, 2016). Under the rearing conditions reported here, the larval to adult stage of *An. coustani* decreased across the generations and exhibit a different pattern of adult emergence (Figure 3). Indeed, F1 adults emerged in two peaks. The first peak occured between day 16 to day 18 from the day of egg laying and the second peak at day 20. From F2 to F4 adult emergence follows a 2 peak pattern, but each peak occurs earlier, except the second peak of F2 that was at day 20 as for the F1. Interestingly, F5 exhibited a broad single peak of adult emergence between day 11 and day 17, while F6 a sharp single peak of adult emergence centered at d13. These changes in adult emergence pattern possibly reveals a trait of adaptation to colonization.

**Figure 3:**
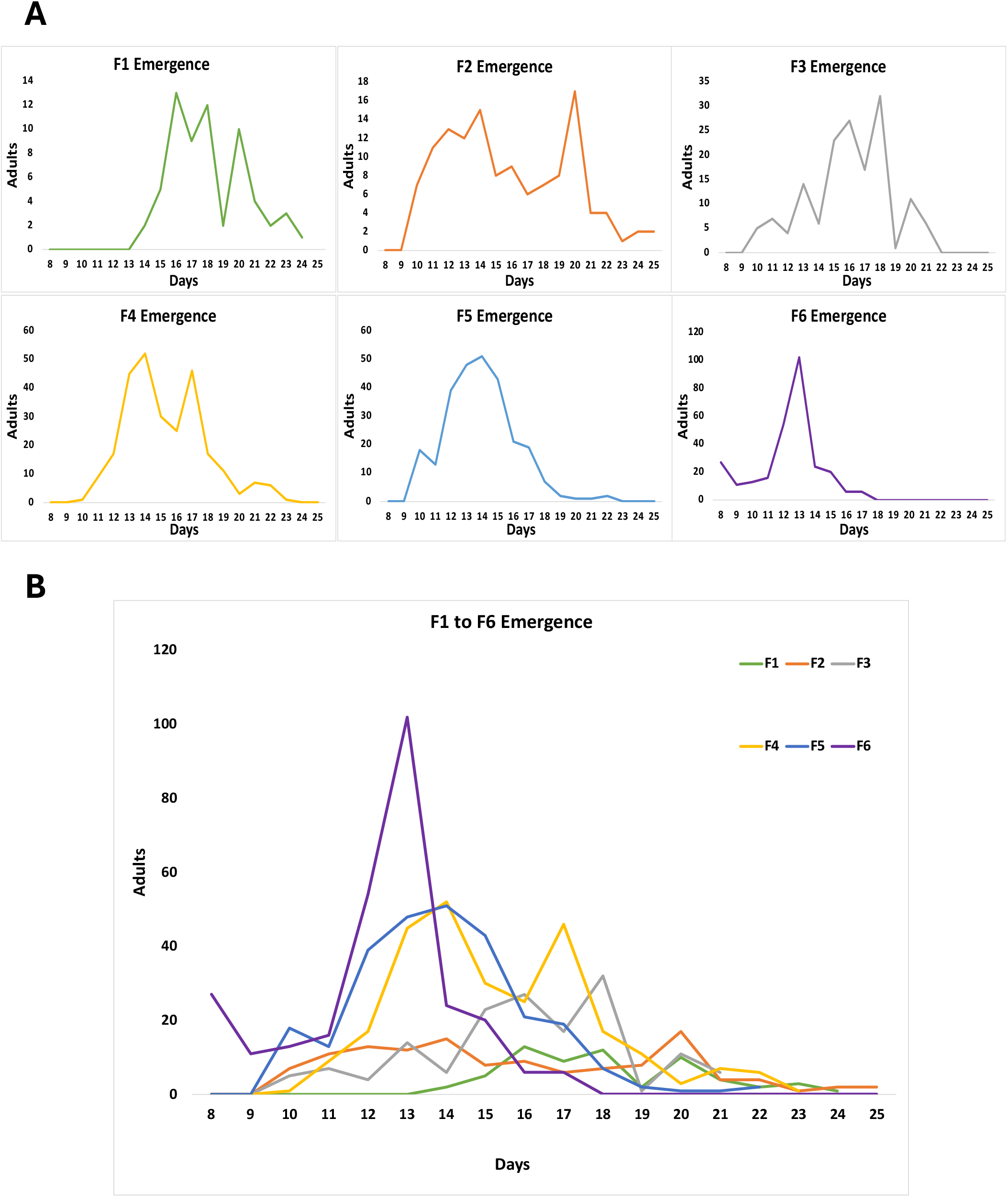
Graphic representation of *An. coustani* adult emergence at each generation. A) individual graphs showing the pattern of emergence of F1 to F6. B) Aggregated graphs highlighting the clear changing patterns of adult emergence from F1 to F6 both in mode and date of the peak of emergence. The scale of the abscissa axis corresponds to days till the first appearance of L1.

## Concluding remarks

We report here the first successful estabishment of a colony of *An*.*coustani*. As this mosquito species would not mate in cage, each of the 8 generations produced resulted from forced mating. This is comparable to the *Anopheles cracens* colony recently established to the 6^th^ generation by forced mating [1]. On the contrary, a novel *Anopheles atroparvus* colony was established to the 10^th^ generation without forced mating [5]. *An. coustani* and *An. atroparvus* being from the *Anopheles* subgenus, while *An. cracens* belongs to the *Cellia* subgenus no conclusion can be drawn as whether species from the *Anopheles* subgenus would required forced copulation for colony establishment. Indeed, even a reference colony of *Anopheles dirus* (*Cellia* subgenus) is maintained by forced copulation at the MR4 repository (WRAIR2 strain, https://www.beiresources.org/Catalog/BEIVectors/MRA-700.aspx)

As maintaining a colony by forced copulation is highly demanding in human resources and skill, the biggest refinement would be to obtain a free mating colony. Towards this goal, it would be of great interest to test whether the tools developed for establishing *An. darlingi* colonies would be beneficial, notably the use of light flashes to stimulate free copulation and shift in night temperature [2, 30, 37]. As discussed previously, additional refinements in the larval feeding regimen and larval density would possibly contribute to decrease the mortality rate during both larval and nymph stages.

Overall, having at hand an *An. coustani* colony whether free mating or maintained by forced copulation as *An. dirus* will provide an unvaluable tool to better understand the biology of this secondary malaria vector and assessing relevant malaria transmission control strategies.

## Acknowledgements

We wish to thank the technicians of the Medical Unit from Institut Pasteur de Madagascar for their support in the field and the insectary work (Tata, Lala, Fidélis, Andrisoa, Onja). Thanks as well to Jean Pierre Ratavilahy, our guide in Andriba ; to Dadabe, the owner of the zebu’s park ; and members of the CSBII and of the city hall in Andriba. This work was supported by funds from Institut Pasteur de Madagascar to RG and a Girard PhD grant to TMA who also received an award grant from L’Oréal-Unesco-FWIS Jeunes talents Sub-saharienne. The funders had no role in study design, data collection and analysis, decision to publish, or preparation of the manuscript.

## Author Contribution

Conceived the project : TMA, CB and RG. Designed the experiments: TMA, NP, CB. Performed the experiments: TMA, FTR, MRA, MRR and HJVS. Analyzed the data: TMA and CB. Drafted the paper: TMA and CB. Reviewed and edited the manuscript: TMA, RG and CB. All authors contributed to the article and approved the submitted version.

